# Computational Model for Biological Neural Network for Localisation of Sound in the Vertical Plane

**DOI:** 10.1101/2020.08.05.238402

**Authors:** Anandita De, Daniel Cox

**Affiliations:** Department of Physics, UC Davis

## Abstract

We build a computational rate model for a biological neural network found in mammals that is thought to be important in the localisation of the sound in the vertical plane. We find the response of neurons in the brain stem that participate in the localisation neural circuit to pure tones, broad band noise and notched noise and compare them to experimentally obtained response of these neurons. Our model is able to reproduce the sensitivity of these neurons in the brain stem to spectral properties of sounds that are important in localisation. This is the first rate based population model that elucidates all the response properties of the neurons in the vertical localisation pathway to our knowledge.

## 1 Introduction

The interaction of sound with the outer ear (pinna) modifies the energy in certain frequency bands in the spectrum of a broadband noise in a way dependent on the location of the sound in the vertical plane.The function mapping the original sound spectrum to the modified one after interaction with the outer ear is known as the head related transfer function (HRTF) [1], [2], [3]. More specifically, the interaction of the sound with the outer ear produces notches (frequency bands where energy is decreased) in the spectrum; the frequency and magnitude of these notches are a function of the angle of elevation of the sound source.

There is evidence that there are neurons in the auditory pathway dedicated to the processing of these notches. These neurons are located in the dorsal cochlear nucleus (DCN) and the inferior colliculus (IC) and are sensitive to the frequency and the magnitude of the notches. Therefore they are believed to be important in the localisation of sound in the vertical plane.

Fig 1 shows the neural circuit that has been proposed to describe the response of the type 4 neurons of the DCN and the type O neurons of the IC [4], [5]. In this paper, we build a firing rate model at steady state for the neural circuit proposed above. This is a feed forward neural network with each layer consisting of the neurons which are relevant for the neural circuit which process these notches. Our model describes how neurons up to the pre-cortex area are tuned to spectral notches essential for the localisation of sound in the vertical plane.

**Figure 1.**
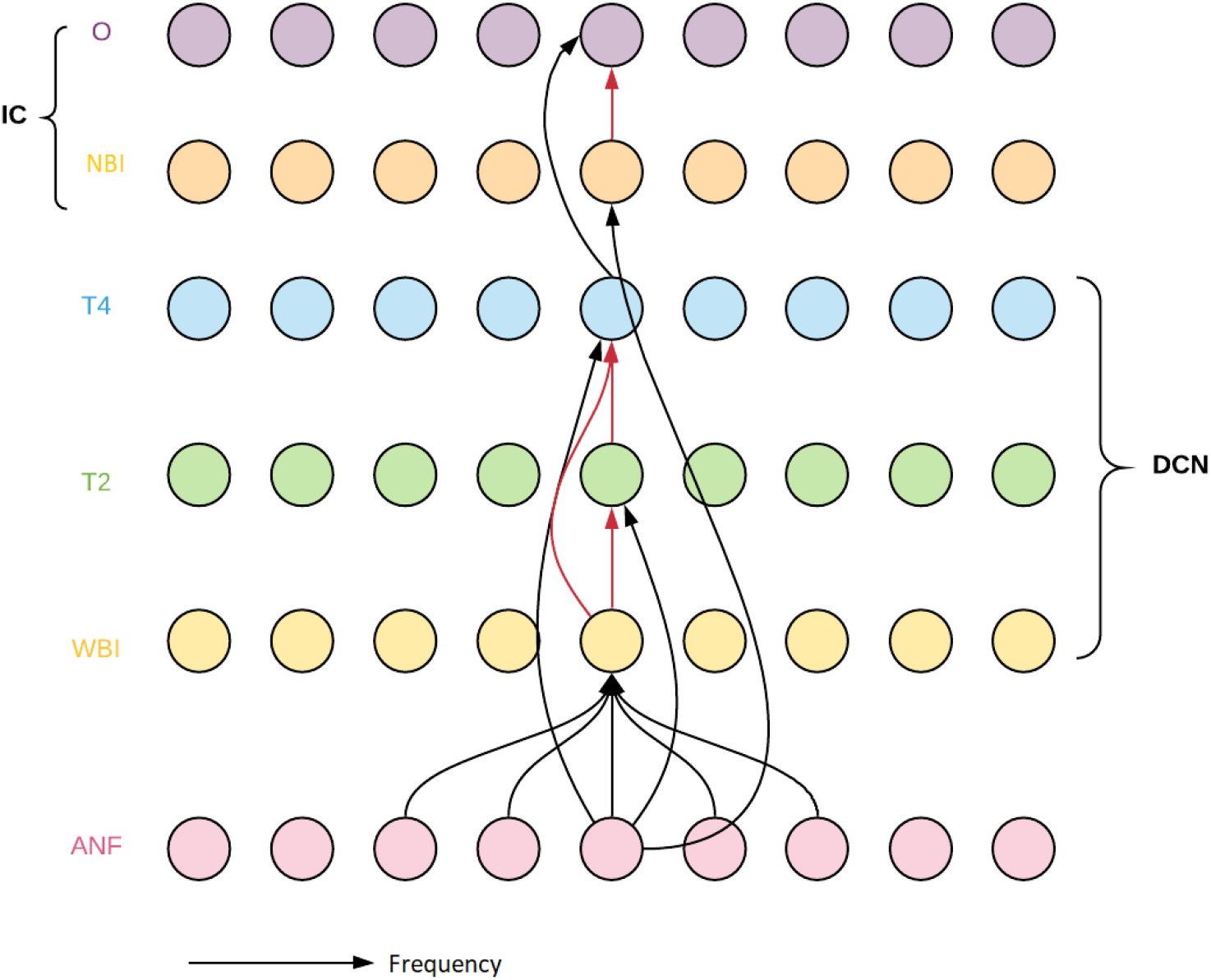
Model Schematics. Schematic of feedforward neural network with 6 layers. Each layer is labelled on the left. Black arrows represent excitatory connections and red arrows represent inhibitory connections. The arrows only represent the connections, the actual weight matrices are given in 5.

We consider the response of the relevant neurons to three kinds of stimuli that have been experimentally recorded : pure tones, broad band noise and notched noise which is a simulation of the notch produced by the HRTF on the spectrum of the sound. The different stimuli that we will be considering for our model are: a) pure tone at a Best Frequency (BF), b) pure tone frequencies swept across the entire range of BFs at different intensity levels, c) broadband noise at different spectrum levels d) Notched noise at different spectral intensity levels, e) notch center frequencies swept across the entire range. The best frequency(BF) of a neuron is defined as the frequency of pure tone for which the neuron responds above its spontaneous rate at the lowest intensity. For a broad band noise we consider a white noise with equal power in each of the frequency components with a certain center and width. We are of course most interested in the notched noise response of the type 4 neurons and the type O neurons since these are sensitive to the frequency of the notches. The complex response properties of these neurons to the above mentioned stimuli show the non trivial integration properties of these neurons which makes the modelling task a challenge.

We consider a firing rate model at steady state. One of the reasons for this is that the experimental results that we compare our results with are rate vs level functions for the different neurons or rate vs best frequency of the neurons. Since we do not have experimental data on rate as a temporal function for the different neurons we are considering, we forego it at our first pass for modelling this circuit. The second reason for considering a model at steady state is because we are interested in the interactions between the different populations of neurons in our model and how it gives rise to complex response properties.

Our motivation in developing this model is to see how the localization network responds when damaged by a neurodegenerative disease such as Alzheimer’s, which is known to impair the ability of afflicted individuals to localize sound, particularly in crowded environments [7]. Moreover, there is substantial evidence to suggest that the auditory brain stem itself is directly impacted by Alzheimer’s driven neurodegeneration [8].

We show that our model is able to reproduce the responses of the neurons recorded in the DCN and the IC at least qualitatively. Though we are not able to capture all the features of the experimentally recorded neurons, we are able to reproduce the features that are important in the localisation of sound. In section 2, we describe our model in more details including the behaviour of the neurons in the different layers and the connections between the layers. In section 3, we describe the successes and the shortcomings of our model. In section 4, we discuss the novel features of our model.

We have built a rate based population model to explain the responses of the type 4 neurons and the type O neurons. Our model architecture has been inspired by the model by Blum and Reed [10], [12]. However, Blum and Reed did not construct a model with neurons logarithmically arranged in frequency as the auditory fibers are known to be. Their neurons were also not correlated with observed best frequencies. Additionally they did not consider responses to notched noise sweeps and their model did not extend to the type O neurons which have been discovered more recently. The arrangement of the neurons according to their BFs and the integration of inputs over some range of frequencies around the BF was inspired by the models of Hancock and Voight [13], [6]. However, the models of Hancock and Voight are spiking models and hence it is more difficult to simulate and study the effects of integration of the inputs between different kinds of neurons. Our model is different from that of [14] in that it is a population model and involves interactions between populations of neurons and has more realistic weight functions connecting populations of neurons.

## 2 The Model

The model is a feed forward neural network consisting of six layers of neurons. Each layer has a hundred neurons that are tonotopically arranged, ie, they are arranged according to the frequency to which they are most sensitive. In our model, we have the BFs of the neurons spanning four octaves, starting from 1.25 kHz and going up to 20 kHz with 25 neurons in each octave. So the BF of each successive neuron is 2^0.04^ times the BF of the previous neuron. This is a reasonable hearing range for humans. We refer to all the neurons below by their BFs.

It is experimentally known that the tonotopic arrangement of the neurons are preserved in the higher areas of the auditory pathway, so we have arranged our model in this way, which was also done in [6] and [10]. However in [10], the authors did not map their frequencies into the actual range of hearing frequencies like we do. Our model is also easily scalable in the number of neurons. Increasing the number of neurons will increase the number of neurons per octave and decrease the difference in the BFs.

The first layer consists of the auditory nerve fibers which receive all auditory stimuli. The auditory nerve fibers transmit this stimuli to the DCN which is the first relay station in the auditory pathway. The neurons in the second, third and the fourth layer in our model lie in the DCN. The neurons in the DCN project to neurons in the IC. The neurons in the fifth and sixth layer of our model lie in the IC. We will describe the neurons in each of these layers and their experimentally recorded response properties in details below.

### Auditory Nerve Fibers

In this section, we define the rate function to model the response of an auditory nerve fiber for pure tone stimulus based on its receptive field. Then we generalise it to a response for more complicated stimulus like broad band noise. The auditory nerve fibers innervate the hair cells which are present on the basilar membrane. The basilar membrane is a thin coiled membrane with a gradient in tension and thickness along its axis. So the different regions of the basilar membrane have different resonant frequencies. The auditory nerve fibers connected to a particular region the basilar membrane is most sensitive to the resonant frequency of that region. The auditory nerve fibers are tonotopically arranged because they connect to different regions of the basilar membrane.

In neuroscience literature [11], the rate of a neuron as a function of a stimulus parameter *s*, *r* = *f* (*s*), is defined as a tuning curve. By this definition, (1) would be called the tuning curve of the ANFs for a pure tone. The tuning curve of an ANF is defined as the minimum intensity as a function of pure tone frequency that elicits a response from the ANF above its spontaneous rate. We first define the rate of an ANF in the first sense, ie as a function of stimulus parameters, the frequency of a pure tone *f_PT_*, and the intensity of a pure tone *s*.

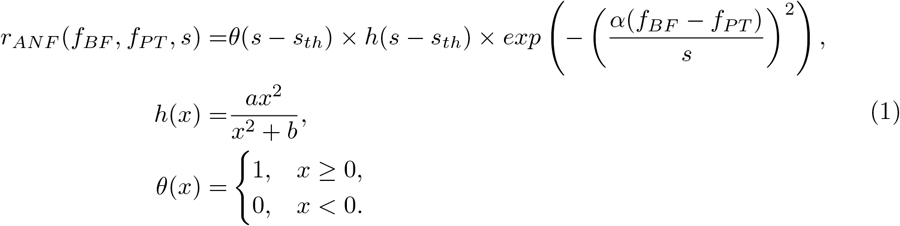

This equation is inspired by tuning curves for other neurons found in the literature. This is the first such function to be defined for an ANF to our knowledge. All other models have only considered rate as a function of intensity, which is the second factor on the right side of (1). This was modelled based on the observation that as the intensity increases, the frequency range over which the pure tone elicits a response from an ANF also increases.The first figure in Fig 2, shows the rate of an ANF as a function of the frequency of pure tones at different intensities according to (1). The second figure in Fig 2, shows the saturating function *h*(*x*) for different parameters. Table 1 gives the values of the parameters in (1) and their description.

**Table 1.** Parameters in (1)

| Parameters | Function of the Parameter |
| --- | --- |
| $s_{th}$ | Threshold for ANF (20 dB) |
| $a$ | Determines the value of saturation of firing rate (200) |
| $b$ | Determines the slope of the saturating function $h$ |
| $\alpha$ | Determines the width of the tuning curve Fig 2 (a) |

**Figure 2.**
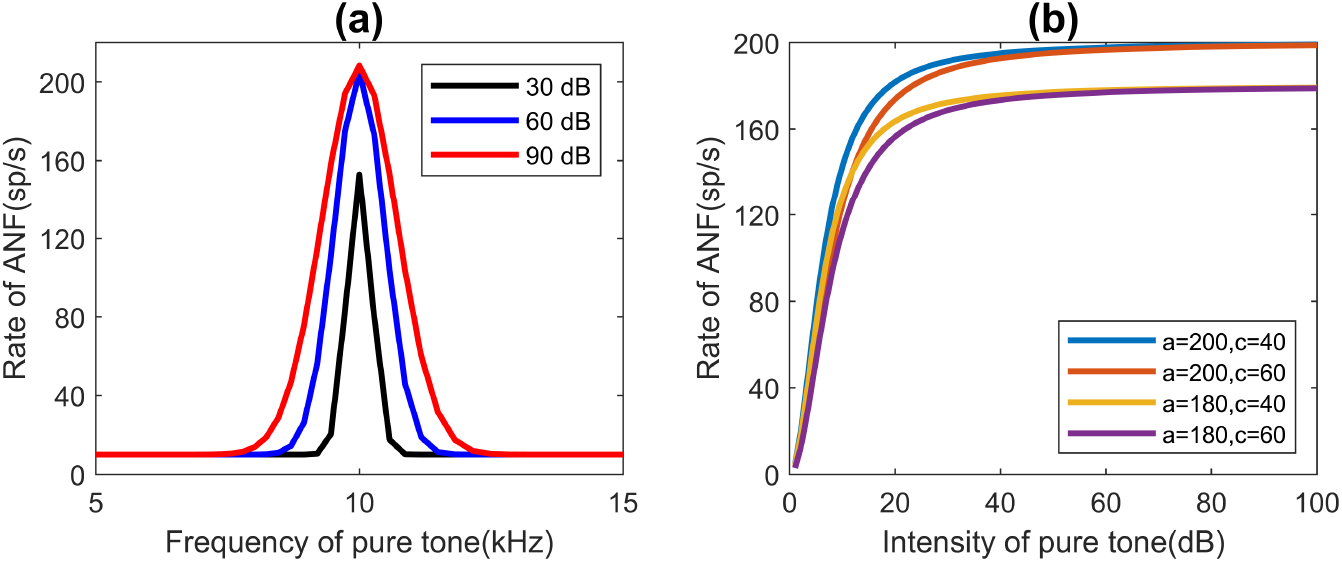
Response of Auditory Nerve Fibers (ANF) as a function of frequency and intensity. a) Rate vs frequency of pure tones for an ANF(BF=10 kHz) at different intensities (*α* = 90) obtained from (1). b) The saturating function with different parameters.

On the left of Fig 3 we have the tuning curves obtained experimentally. On the right of Fig 3 we have the tuning curves obtained from (1). To get the tuning curves *s*(*f_BF_, f_PT_*) from (1), we set *r_ANF_* (*s, f_PT_, f_BF_*) = *s_th_*. Since the function (1) is symmetric about *f_BF_* for the pure tone frequencies we do not get the long tail extending to the lower frequencies at higher intensities as is seen experimentally. We would need to modify (1) to include this feature.

**Figure 3.**
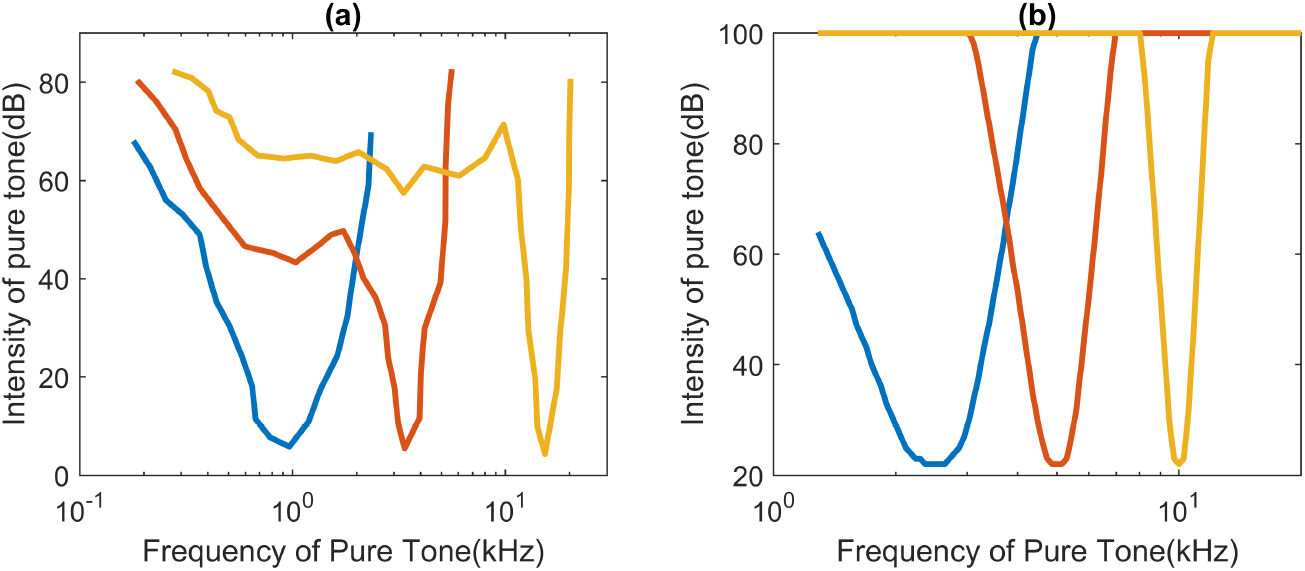
Comparison of experimental and model tuning curves for ANFs. a) Tuning curves for ANFs obtained experimentally. The tuning curve for an ANF is intensity as a function of pure tone *s*(*f_pt_*), at which the ANF responds above its spontaneous rate. Adapted from ([9]) b) Tuning curves obtained from the model of the ANFs above.

We model a broadband noise to have a constant flat spectrum over a range of frequencies. We assume that the response of an ANF to such a stimuli is a saturating function of the power in the frequency component corresponding to the BF of the ANF. So the ANFs whose BFs lie under the nonzero power spectrum of the noise respond to the noise and the others do not.The first figure in Fig 4 shows the noise stimulus and the second one shows the response of the ANFs to the broadband noise on the left. Similarly Fig 5 shows a notched noise stimulus on the left and the rate of the ANFs vs the BFs of the ANFs for the stimulus to the right.

**Figure 4.**
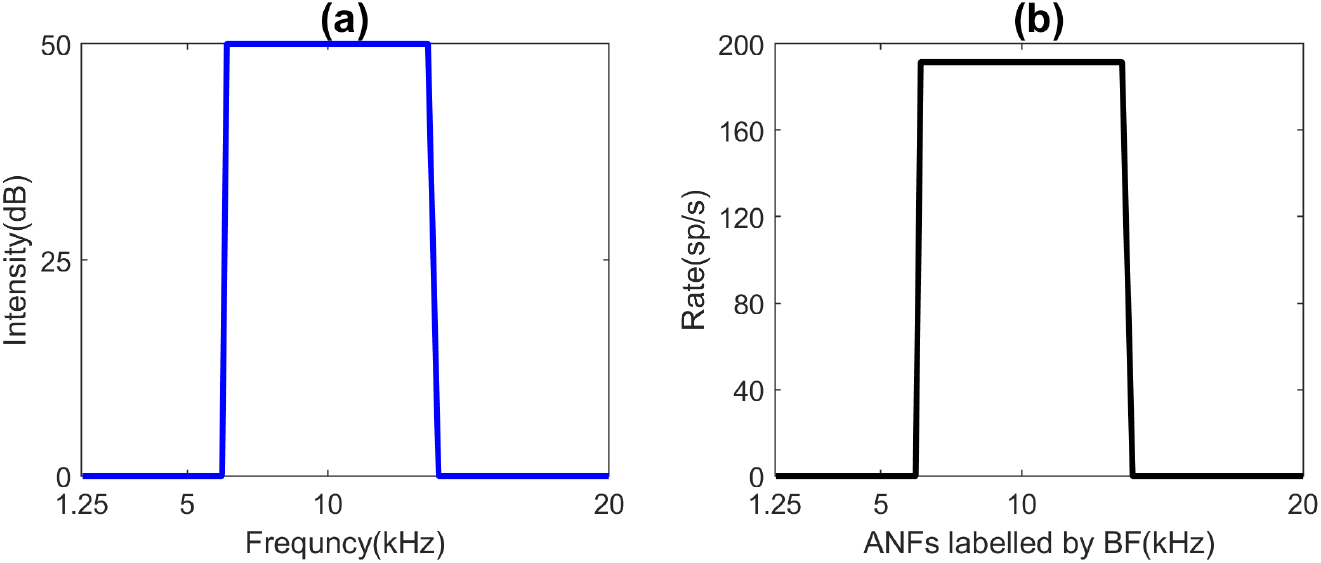
Response of ANFs to broadband noise. a) Broadband noise with a constant power spectrum of 50 dB centered on 5kHz having a width of 7kHz. b) Response of ANFs labelled by their BFs to the broadband noise on the left.

**Figure 5.**
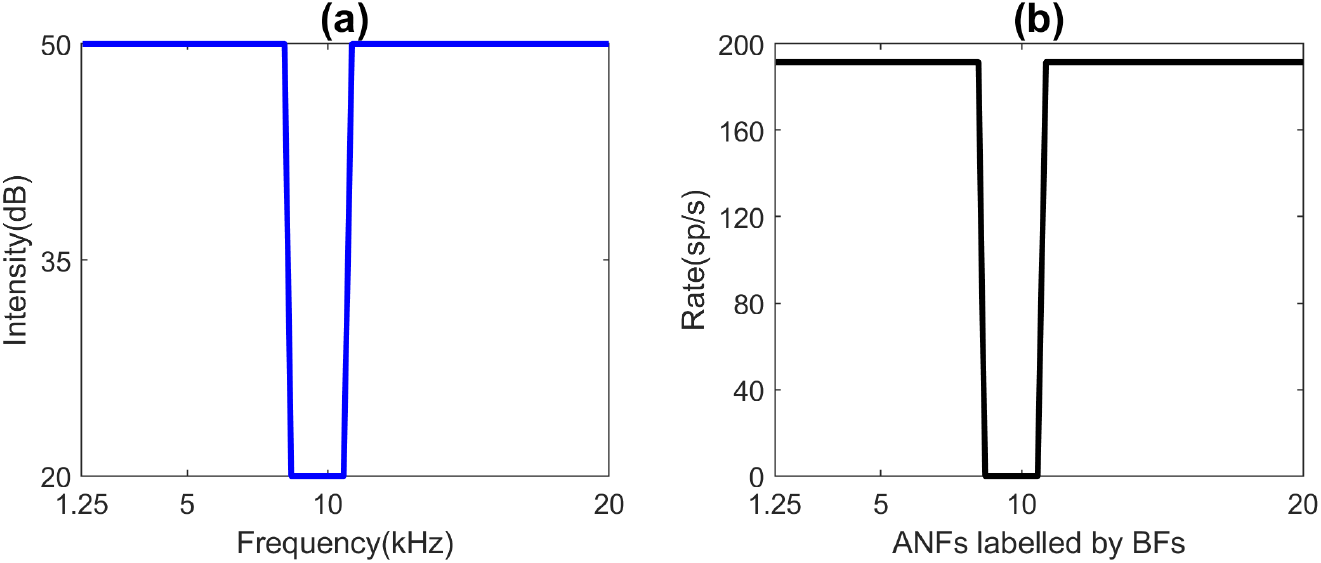
Response of ANFs to notched noise. a) Notch of width 1.6 kHz centered at 9 kHz in a broadband noise of 14 kHz. b) Rate of ANFs labelled by their BFs for the notched noise described in (a).

### Neurons in the DCN

The neurons in the DCN and the connections between them are shown in Fig 1(a). This neural circuit was proposed in [4] to explain the responses of the type 4 neurons of the DCN. They conjectured the existence of the neurons which they called the WBIs which receive weak inhibitory inputs from a wide range of ANF frequencies. So they respond weakly to pure tones and strongly to broadband noise. They showed that the responses of the type 4 neurons and the type 2 neurons could be explained if one considered the inhibition from these cells. The onset chopper neurons in the Ventral Cochlear Nucleus(VCN) are known to have responses similar to that of the WBIs [22].

The type 2 neurons are excited by the ANFs and inhibited by the WBIs. The response properties of these neurons as a result of the interaction of the above two kinds of neurons are discussed in the next section. The type 4 neurons are are excited by the ANFs and inhibited by the type 2 neurons and the WBIs.

We give the input output functions of the above neurons and the weight matrices connecting them in 5. The input output functions are the same as that in [10] but we do not have any justification for these except that they work well. The weight functions that we use are closely related to how receptive fields of neurons are modelled. This kind of weight functions have not been previously used to model this neural network. Our model helps shed light on the receptive fields of the different neurons and their frequency integration properties.

### Neurons in the IC

Davis et al [5] have recorded the response of the neurons in the inferior colliculus to pure tone sweeps and notched noise sweeps. They show that the type O neurons of the IC are sensitive to the position of the notches. They have shown that these neurons receive direct excitatory inputs from the type 4 neurons of the DCN. They also propose a local circuit in the IC to explain the response of the type O neurons to the different stimuli described above. They conjecture the existence of narrow band inhibitors which inhibit the type O neurons and are excited by frequencies in a narrow range lying below the BF of the type O neuron which it inhibits. They call these neurons the Narrow Band Inhibitors(NBI). They also conjectured the existence of neurons which receive input from a wide range of frequencies and excite the type O neurons called the Wide Band Excitators (WBE). For our model, we only consider the NBIs which are necessary to reproduce the sensitivity of the type O neurons to notched noises. We will describe below what we fail to reproduce as a result of not considering the WBEs. It has been conjectured in [5] that the type I and type V units in the IC might be the narrow band inhibitors and the onset units in the IC could be the WBEs but more experiments are needed to confirm these conjectures.

## 3 Results

The type 2 neurons respond strongly to pure tones at their BF and are characterised by their non-monotonic rate level functions. Fig 6(a) shows the rate level function obtained from our model. We get a non-monotonic rate level function as predicted by experiments. This is because, as intensity increases, the range of ANFs which respond to a pure tone increases. Therefore the rate of the WBIs as the intensity of the pure tone increases. Since the WBIs inhibit the type 2 neurons, the increase in their rate brings down the rate of the type 2 neurons at higher intensities of pure tones. This lends some support to the widening of the response curves of the ANFs as the intensity increases and to the existence of WBIs. Fig 6(b) shows that the overall rate of the type 2 neurons decreases as the width of the broadband noise increases. This can again be explained by the fact that the response of the WBIs increases as the width of the broadband noise increases. Type 2 neurons are also experimentally found to have a high threshold which leads to the neurons being completely shut off if the input is below the threshold. In our simulations the type 2 neurons are completely shut off by a notched noise when the notch lies above its BF.

**Figure 6.**
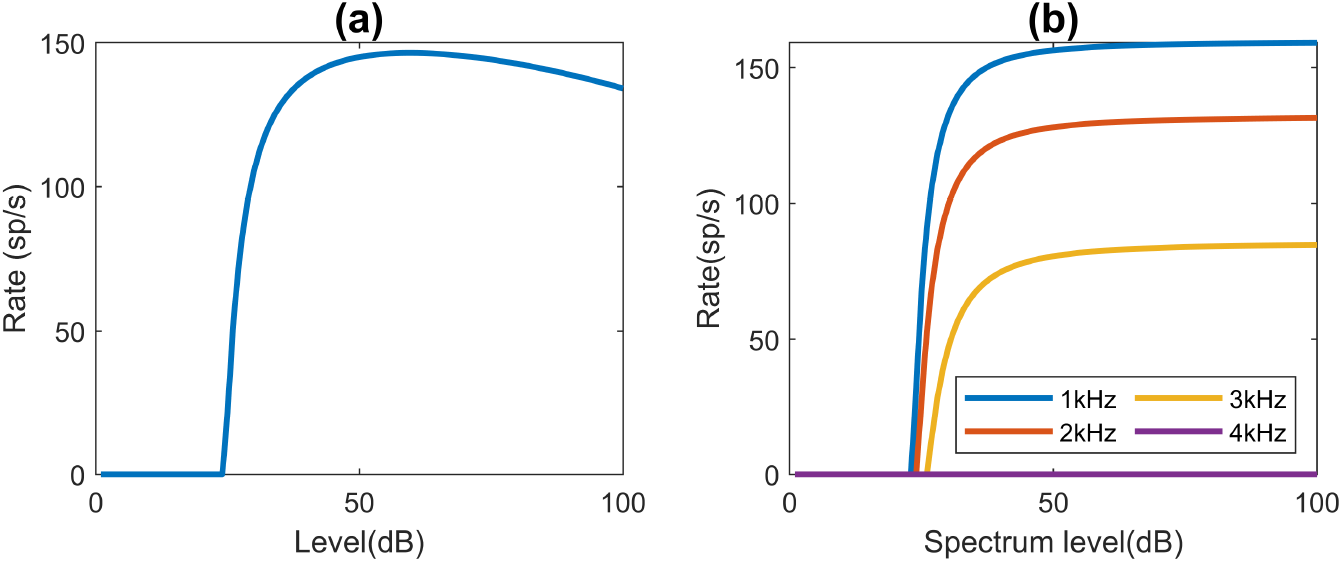
Response of type 2 neurons. a) Rate vs level curve for the response of a type 4 neuron to a pure tone at BF. b) Rate vs level curve of a type 2 neuron for noise. We can see that the rate decreases as the width of the broadband noise increases which is also observed experimentally.

The type 4 neurons are excited for a narrow range of frequencies at low levels(dB) and are inhibited or shut off for higher intensities. Fig 7(a), shows the rate vs level curve for a pure tone at its BF for a type 4 neuron. The type 4 neurons are modelled such that they have a spontaneous rate of 30 kHz. We see that the rate increases above the spontaneous rate as intensity increases above 20 dB which is the threshold for the ANFs. At low intensities the type 4 neurons only receive excitatory inputs from the ANFs since the type 2 neurons have a high threshold as we discussed above and the WBIs respond very weakly at low intensities. As the intensity increases, the inhibition of the type 2 neurons and the WBIs soon overcome the excitation from the ANFs. So we have a narrow peak in the rate vs level curve for the type 4 neurons for pure tones. Fig 7(b) shows the rate vs spectrum level of broadband noise for type 4 neurons. The type 4 neurons are excited above their spontaneous rates by broadband noise of different widths. The rate goes down as the width of the broadband noise increases because the inhibitory inputs from the WBIs increase.

**Figure 7.**
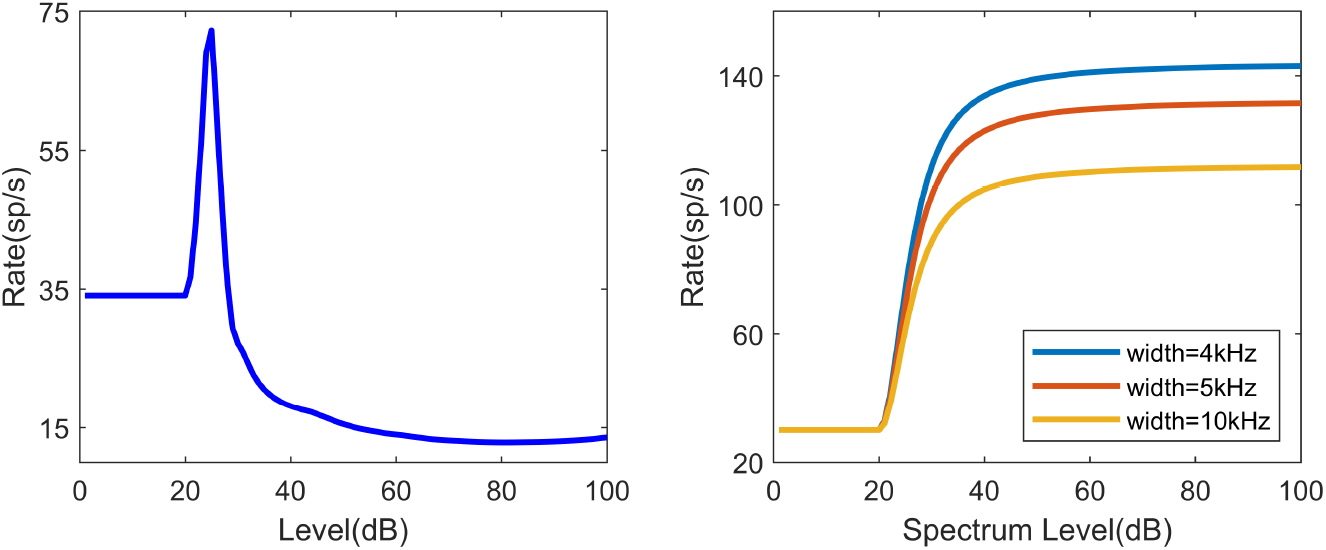
Response of a single type 4 neuron to pure tone at BF and broad-band noise. a) Rate vs level curve for a type 4 neuron to a pure tone at its BF. It is excited above its spontaneous rate in a narrow frequency range above the threshold and it is inhibited for higher decibels. b) Rate vs spectrum level curves of a type 4 neuron for broadband noise.

Fig 8(a) shows the response of the type 4 neurons for pure tone sweeps at different levels. As we discussed above, the type 4 neurons are excited above their spontaneous rates for a narrow range of frequencies at low levels. At higher levels, the rate of the type 4 neurons fall below the spontaneous rate. We failed to reproduce the inhibition of the type 4 neurons for pure tones in the entire frequency range for higher levels. There is inhibition only over the range from which the WBIs receive inputs. Fig 8(b) shows the response of a type 4 neuron with BF 9.2 kHz to notched noise sweeps. The notch has a width of 1.6 kHz and its center is swept over 3 octaves. The type 4 neurons are inhibited below their spontaneous rate when the notch lies over the best frequency of the neuron and they are excited above their spontaneous rate when the notch center moves away from the best frequency. We have not been able to reproduce the different ranges of inhibitions of the neurons at different spectrum levels of the notched noise. Experiments have shown that there is maximum inhibition at intermediate levels.

**Figure 8.**
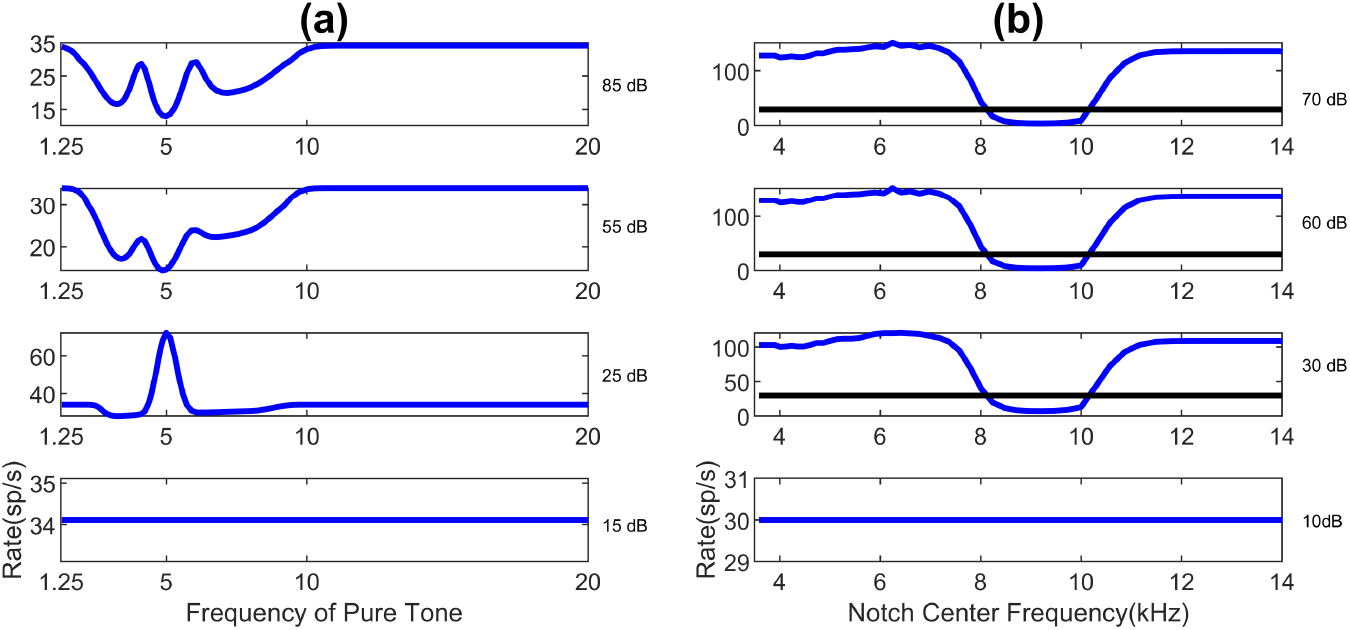
Responses of Type 4 neurons to pure tone sweeps and notched signal sweeps. a)Rate vs frequency of pure tone of a type 4 neuron with BF 5 kHz for pure tone sweeps at different levels. b) Rate vs notch center frequency of a notched noise with notch width 1.6 kHz which is swept over 3 octaves, of a type 4 neuron with BF 9.2 kHz.

Fig 9(a) shows the rate vs pure tone frequency curve for a type O neuron with BF 5 kHz at different levels. The type O neurons are excited above spontaneous rates at low levels and a narrow range of pure tone frequencies. This behaviour is similar to the type 4 neurons of the DCN. The drop in the rates just below the BF is due to the inhibition from the NBIs whose BFs lie in a narrow range of frequencies below the BF of the type O neuron which they inhibit. As the intensity increases the rates of the type 4 neurons themselves fall below spontaneous rates and hence the type O neurons are no longer excited above their spontaneous rates either. Fig 9(b) shows the rate vs notch center frequency curves for a type O neuron with BF 9.2 kHz at different spectrum levels of the notched noise. The type O neuron is excited above its spontaneous rate when the notch lies just below the BF of the neuron. This is because in our model the type O neurons are inhibited by NBIs whose BFs lie below the BF of the type O neuron it inhibits. So when the notch lies over the BFs of the NBIs, the type O neurons do not receive any inhibition and are excited by the type 4 neurons. The width of the excitation depends on the integration of inputs from the type 4 neurons and the NBIs.

**Figure 9.**
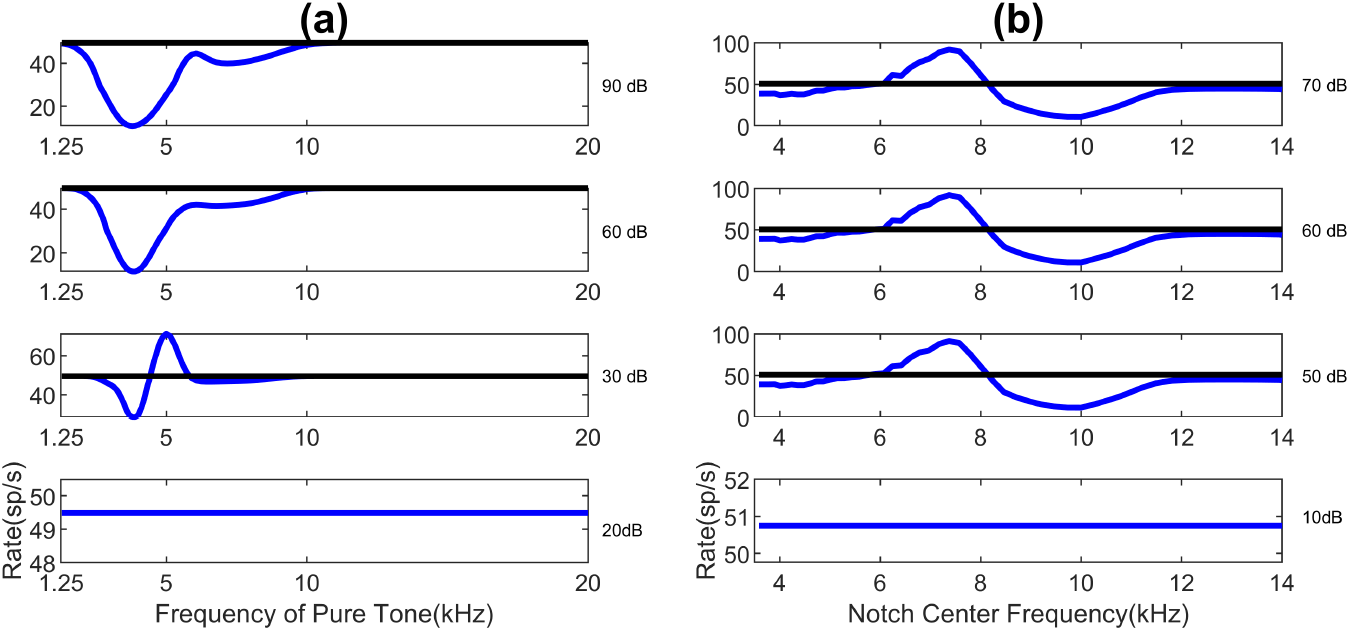
Response of type O neurons to pure tone sweeps and notched noise sweeps at different levels. The black lines represent the spontaneous rates. a) Rate vs pure tone frequency of a type O neuron with BF 5 kHz. The neuron is excited for a narrow range of pure tone frequencies at a low intensity like type 4 neurons. b) Rate vs notch center frequency of a type O neuron with BF 9.2 KHz. The notch had a width of 1.6 kHz and was swept across a range of 2 octaves from 3.5 kHz to 14 kHz.

## 4 Discussion

We have been able to reproduce the main features of the response of the type 4 neurons and the type O neurons to notched noises and shown their sensitivity to the frequency of notches as found in experiments. The type 4 neurons of the DCN have been experimentally found to be excited by a narrow range of pure tone frequencies at BF at low intensities and inhibited at high intensities. They are inhibited by notched noise when the notch lies over the BF of the neuron. The width of the inhibitory regions for both the pure tone and the notched noise do not match the experimental results but we have reproduced the two main features characterising the type 4 neuron mentioned above as shown in Fig 8.

We have also reproduced the general characteristics for the response of the type O neurons as shown in Fig 9. The pure tone response characteristics are the same as that of the type 4 neurons of the DCN where they have a narrow region of excitation at low intensities and are inhibited at high intensities. However, unlike the type O neurons these neurons are excited when the notch center lies below their BF. We have not been able to reproduce the broad excitation of the type O neurons at low intensities. This could be because we have not included the wideband excitators that were conjectured in [5].

We use an approximation for the responses of the auditory nerve fibers for complex stimuli like broadband noise and notched noise. The response of the auditory nerve fibers are crucial since it it is the first layer and all subsequent layers receive inputs from the ANFs, which might be one of the reasons that some of the response features to complex stimuli for the type 4 neurons and type O neurons are missing. More accurate responses for the ANFs have to be modeled by filter functions on the stimuli [16], [17] which are in the time domain. For this work we have assumed our neurons to be in steady state. To match experimental results more closely, we would have to simulate a dynamical model for all our neurons and Fourier transform it to obtain rate vs frequency curves. Even so, rate vs intensity curves would be difficult to obtain from such models. We leave the development of such a dynamical model to future work. As we have mentioned earlier, the parameters in the weight matrices of our model might not be optimal since they were found by a trial and error method. Optimising these parameters would also bring the responses closer to their experimental analogs.

Recently, convolutional neural nets (CNNs) have been used to model retinal responses to natural stimuli and to determine the receptive fields of neurons in the visual pathway [15]. This neural network can also be thought of as a CNN with a single filter (a frequency dependent filter). It could be trained on various natural stimuli with experimentally found responses to find the receptive fields of the neurons. We could then compare the proposed receptive fields to the receptive fields obtained by such training. In a different approach, we could train our neural network on natural stimuli with experimentally obtained responses to optimise the parameters in the weight functions defined below. This would be a more algorithmic approach to determine these parameters.

As the model stands now, we have shown that the neurons in the DCN and IC are important in the processing of notches. A full mapping between the position of these notches to the vertical location of the sound has not yet been found. It is yet to be experimentally found how the inputs from the IC are processed in the auditory cortex. The final aim of this neural circuit would be to have a mapping between the position of the notch in the spectrum to the location of the source of the sound.

## Acknowledgements

We would like to thank Mitch Sutter for a discussion on the localisation of sound at the beginning of this work. AD would like to thank Mark Goldman and his group for inputs during group meeting discussion.

## 5 Supplementary Information

### Mathematical Description of the Model

In this section we give the equations for our feed forward neural network. As has been already discussed, we are considering a rate model at its fixed point. This can be easily generalised to a dynamical model by looking at the stimuli as a function of time and then performing Fourier transforms to obtain rate vs frequency plots. The different weights that we use are given in Fig 10

**Figure 10.**
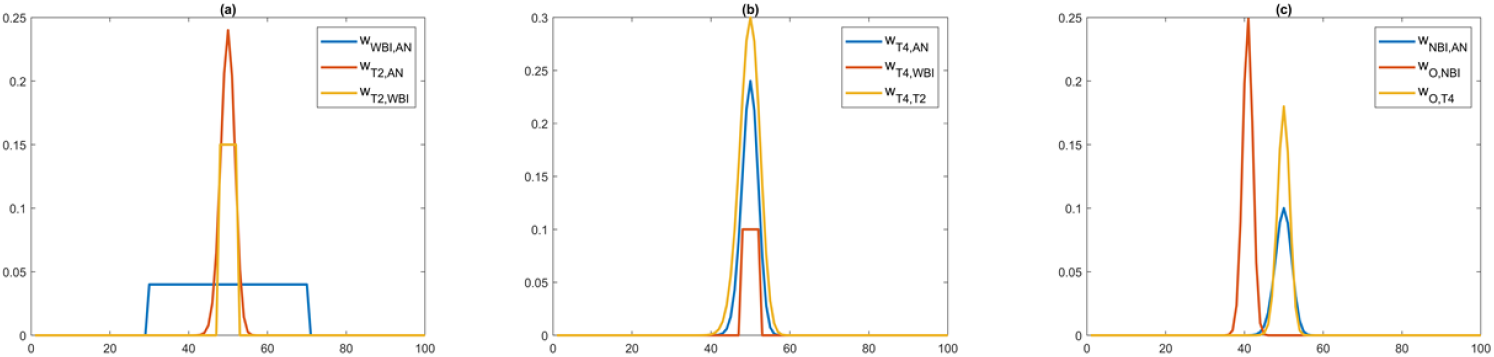
Weights connecting the different neurons in our model. a) Weights from ANFs to WBI(*blue*) and weights from WBI and ANFs to T2 with BF 10 kHz. (b) Weights from ANF, WBI, T2 to T4 neuron with BF 10 kHz. (c)Weights from ANF to NBI(*blue*) and weights from NBI, T4 to O.

The type 2 neurons receive excitatory inputs from the ANFs and inhibitory connections from the WBIs.

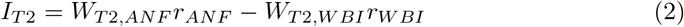

*f* (*x*) gives the input output function for ANFs. This input output function was taken from [10]

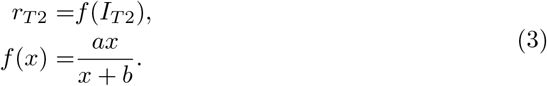

In our model, *a* = 250 *sp/s, b* = 70 *sp/s* and the T2 neurons have a threshold of 35*sp/s*. Note that the units of the parameter *a* and *b* are such that *r*_*T*2_ has the units of spikes/s.

Type 4 neurons receive excitatory connections from ANFs and inhibitory connections from type 2 neurons and WBIs.

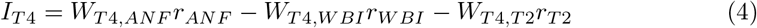

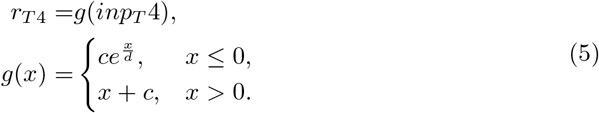

For our model, *c* = 30*, d* = 30.

The type O neurons of the IC are excited by the type 4 neurons of the DCN and inhibited by the narrow band inhibitor(NBI) in the IC.

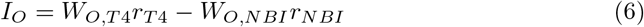

The input output function for type O units were the same as that of the type 4 units.

## Notes

### Competing Interest Statement

The authors have declared no competing interest.

## References

1. May BJ, Huang AY. Spectral cues for sound localization in cats: A model for discharge rate representations in the auditory nerve. The Journal of the Acoustical Society of America. 1997;101(5):2705–2719.

2. Musicant AD, Chan JC, Hind JE. Direction-dependent spectral properties of cat external ear: New data and cross-species comparisons. The Journal of the Acoustical Society of America. 1990;87(2):757–781.

3. Rice JJ, May BJ, Spirou GA, Young ED. Pinna-based spectral cues for sound localization in cat. Hearing research. 1992;58(2):132–152.

4. Nelken I, Young ED. Two separate inhibitory mechanisms shape the responses of dorsal cochlear nucleus type IV units to narrowband and wideband stimuli. Journal of neurophysiology. 1994;71(6):2446–2462.

5. Davis KA, Ramachandran R, May BJ. Auditory processing of spectral cues for sound localization in the inferior colliculus. Journal of the Association for Research in Otolaryngology. 2003;4(2):148–163.

6. Hancock KE, Voigt HF. Wideband inhibition of dorsal cochlear nucleus type IV units in cat: a computational model. Annals of biomedical engineering. 1999;27(1):73–87.

7. Tuwaig M, Savard M, Jutras B, Poirier J, Collins DL, Rosa-Neto P, et al. Deficit in Central Auditory Processing as a Biomarker of Pre-Clinical Alzheimer’s Disease. Journal of Alzheimers Disease. 2017;60(4):1589–1600. doi:10.3233/jad-170545.

8. Mansour Y, Blackburn K, Gonzalez-Gonzalez LO, Calderon-Garciduenas L, Kuleszaa RJ. Auditory Brainstem Dysfunction, Non-Invasive Biomarkers for Early Diagnosis and Monitoring of Alzheimer’s Disease in Young Urban Residents Exposed to Air Pollution. Journal of Alzheimers Disease. 2019;67(4):1147–1155. doi:10.3233/jad-181186.

9. Purves D, Augustine GJ, Fitzpatrick D, et al. editors. Tuning and Timing in the Auditory Nerve. Neuroscience. 2nd edition. Sunderland (MA): Sinauer Associates; 2001. Available from: https://www.ncbi.nlm.nih.gov/books/NBK11105/

10. Blum JJ, Reed MC, Davies JM. A computational model for signal processing by the dorsal cochlear nucleus. II. Responses to broadband and notch noise. The Journal of the Acoustical Society of America. 1995;98(1):181–191.

11. Dayan P, Abbott LF. Theoretical Neuroscience: Computational and Mathematical Modeling of Neural Systems. The MIT Press; 2005.

12. Reed MC, Blum JJ. A computational model for signal processing by the dorsal cochlear nucleus. I. Responses to pure tones. The Journal of the Acoustical Society of America. 1995;97(1):425–438.

13. Hancock KE, Davis KA, Voigt HF. Modeling inhibition of type II units in the dorsal cochlear nucleus. Biological cybernetics. 1997;76(6):419–428.

14. Johnson JS, O’Connor KN, Sutter ML. Segregating two simultaneous sounds in elevation using temporal envelope: Human psychophysics and a physiological model. The Journal of the Acoustical Society of America. 2015;138(1):33–43.

15. Maheswaranathan N, McIntosh L, Kastner DB, Melander J, Brezovec L, Nayebi A, et al. Deep learning models reveal internal structure and diverse computations in the retina under natural scenes. bioRxiv. 2018; p. 340943.

16. Carney LH. A model for the responses of low-frequency auditory-nerve fibers in cat. The Journal of the Acoustical Society of America. 1993;93(1):401–417.

17. Lewicki MS. Efficient coding of natural sounds. Nature neuroscience. 2002;5(4):356–363.

18. Aitkin L, Martin R. Neurons in the inferior colliculus of cats sensitive to sound-source elevation. Hearing research. 1990;50(1-2):97–105.

19. Arle J, Kim D. Simulations of cochlear nucleus neural circuitry: Excitatory–inhibitory response-area types I–IV. The Journal of the Acoustical Society of America. 1991;90(6):3106–3121.

20. May BJ, Huang AY. Sound orientation behavior in cats. I. Localization of broadband noise. The Journal of the Acoustical Society of America. 1996;100(2):1059–1069.

21. May BJ. Role of the dorsal cochlear nucleus in the sound localization behavior of cats. Hearing research. 2000;148(1-2):74–87.

22. Palmer A, Winter I. Best frequency (BF) threshold reductions caused by off-BF non-excitatory tones in onset units of the cochlear nucleus. In: Sixteenth Midwinter Research Meeting of the Association for Research in Otolaryngology; 1993. p. 123.

23. Spirou G, Rice J, Young E. Interneurons of the dorsal cochlear nucleus shape the responses of principal cells to complex sounds. In: Auditory Physiology and Perception. Elsevier; 1992. p. 389–396.

24. Voigt HF, Young ED. Evidence of inhibitory interactions between neurons in dorsal cochlear nucleus. Journal of neurophysiology. 1980;44(1):76–96.

25. Voigt HF, Young ED. Cross-correlation analysis of inhibitory interactions in dorsal cochlear nucleus. Journal of Neurophysiology. 1990;64(5):1590–1610.

26. Young ED, Brownell WE. Responses to tones and noise of single cells in dorsal cochlear nucleus of unanesthetized cats. Journal of Neurophysiology. 1976;39(2):282–300.

27. Young ED, Spirou GA, Rice JJ, Voigt HF. Neural organization and responses to complex stimuli in the dorsal cochlear nucleus. Philosophical Transactions of the Royal Society of London Series B: Biological Sciences. 1992;336(1278):407–413.

28. Young ED, Nelken I, Conley RA. Somatosensory effects on neurons in dorsal cochlear nucleus. Journal of Neurophysiology. 1995;73(2):743–765.

29. Young ED, Rice JJ, Tong SC. Effects of pinna position on head-related transfer functions in the cat. The Journal of the Acoustical Society of America. 1996;99(5):3064–3076.

